# A natural history of networks: Modeling higher-order interactions in geohistorical data

**DOI:** 10.1101/2022.09.26.509538

**Authors:** Alexis Rojas, Anton Holmgren, Magnus Neuman, Daniel Edler, Christopher Blöcker, Martin Rosvall

**Affiliations:** Integrated Science Lab, Department of Physics, Umeå University, SE-901 87 Umeå, Sweden; Gothenburg Global Biodiversity Centre, Box 461, SE-405 30 Gothenburg, Sweden; Department of Biological and Environmental Sciences, University of Gothenburg, Box 461, SE-405 30 Gothenburg, Sweden

**Keywords:** geohistorical data, paleobiology, complex networks, higher-order interactions, Infomap, map equation

## Abstract

Paleobiologists are increasingly employing network-based methods to analyze the complex data retrieved from geohistorical records, including stratigraphic sections, sediments, and fossil collections. However, the lack of a common framework for designing, performing, evaluating, and communicating these studies, leads to issues of reproducibility and communicability. The high-dimensional geohistorical data also raises questions about the limitations of standard network approaches, which assume independent interactions between pairs of components. Higher-order network models better suited for the complex relational structure of the geohistorical data provide an opportunity to overcome these challenges. These models can represent temporal and spatial constraints inherent to the biosedimentary record and describe higher-order interactions, capturing more accurate biogeographical, biostratigraphic, and macroevolutionary patterns. Here we describe how to use the Map Equation framework for designing higher-order network models of geohistorical data, address some practical decisions involved in modeling complex dependencies, and discuss critical methodological and conceptual issues that currently make it difficult to compare results across studies in the growing body of network-based paleobiology research. We illustrate different higher-order network representations and models, including multilayers, hypergraphs, and varying Markov times models, using case studies on gradient analysis, bioregionalization, and macroevolution, and delineate future research directions for current challenges in the emerging field of network paleobiology.

## 1. Introduction: Challenges and opportunities for network-based paleobiology research

Network science is transforming scientific research by enhancing our modeling capacities and helping us understand the behavior of complex systems (Battiston et al. 2020). Many natural and social phenomena can be described as networks, where nodes represent individual components and links indicate their interactions. Network science studies high-dimensional, heterogeneously structured, complex systems and their underlying processes (Barabási and Pósfai 2016). In recent years, standard network models based on pairwise or direct interactions between individual components have been applied to almost every area of paleontological research, including biostratigraphy (Muscente et al. 2019), biogeography (Dunhill et al. 2016; Kiel 2017; Rojas et al. 2017; Kocsis et al. 2018; Jeon et al. 2021), macroecology (Roopnarine 2010), and macroevolution (Muscente et al. 2018; Kocsis et al. 2021). However, methodological inconsistencies and conceptual issues make it challenging to reproduce experiments and compare outcomes across studies, for instance macroevolutionary patterns delineated using standard (Muscente et al. 2018) and higher-order network models (Rojas et al. 2021) representing the same underlying data. The complexity of the high-dimensional and spatiotemporally resolved data retrieved from different geohistorical records, including stratigraphic sections, sediments, fossil collections, and ice cores (National Research Council 2005), also raises questions about the limitations of the standard network models: How accurately do they capture environmental (e.g., sedimentary facies, depositional environments), spatial (e.g., geographical proximity, plate tectonic configuration), and temporal (e.g., chronostratigraphic resolution; geohistorical time arrow) characteristics of the local (bed sets, stratigraphic sections), regional (geological basins; continental records), and global scale systems examined by paleobiologist?

Although network science provides methods for statistical analysis and machine learning of relational data (Brandes et al. 2013; Lambiotte et al. 2019), paleobiologists often describe network analysis as a tool for visualization and qualitative assessment (e.g., Huang et al. 2016; Penn-Clarke and Harper 2020; Ye et al. 2021). This misconception emerges from the general assumption that spatial patterns observed in network diagrams created with force-directed layout algorithms (e.g., Kamada-Kawai algorithm, Fruchterman-Reingold algorithm, etc.) (Csárdi et al. 2024) and representing the observable entities and their connection, properly capture the underlying network dynamics and structure. This assumption ignores the fact that different drawing algorithms highlight different aspects of networks and may not provide reliable insights about their structure. While network visualization techniques are powerful tools for exploratory data analysis (Perri and Scholtes 2020), this misrepresentation reflects a need to adapt research practices in quantitative paleobiology based on theoretical and methodological advances of the network science. Pioneering studies using higher-order representations to explore the deep-time fossil record (Eriksson et al. 2021; Rojas et al. 2021) suggest that the most critical conceptual issue in the growing body of network-based paleobiology research is ignoring the extent to which the choice of network model impacts the results. There are also methodological inconsistencies, including inadequate descriptions of the input network, incomplete explanations of the clustering approach, and uncritical acceptance of the network partition without validation. Overall, the lack of a common framework obstructs interdisciplinarity, reproducibility, and communicability, highlighting the need for standardized research practices in the emergent field of network paleobiology to guide the design, execution, communication, and evaluation of network-based studies.

In this overview, we describe how to use the Map Equation framework (Rosvall and Bergstrom 2008; Edler et al. 2017) to identify important patterns in geohistorical data with different higher-order network models. Specifically, we describe the concept of higher-order interactions in network representations of the fossil record, address some practical decisions involved in modeling the geohistorical data, and illustrate alternative higher-order network models, including multilayer networks, hypergraphs, and varying Markov time models. We illustrate these concepts through classic paleontological examples such as depth gradient analysis, marine bioregionalization, and macroevolution. Our analysis of these case studies either challenges well-established hypotheses or enhances understanding of the underlying research questions, thereby highlighting both the need and opportunity to introduce network models beyond pairwise interactions in paleobiology research. Finally, we delineate major future research directions for current challenges in network paleobiology. We have focused on the Map Equation framework and its applications in paleobiology research because it is an increasingly popular alternative to the standard statistical approaches currently used in paleobiology research (Kocsis et al. 2021; Rojas et al. 2021; Pilotto et al. 2022; Viglietti et al. 2022). The associated software called Infomap for finding community structure in standard and higher-order networks is also widely used in natural and social sciences (Law et al. 2022; Lazaridis et al. 2022; Martins et al. 2022). Overall, our study demonstrates how the Map Equation framework for network analysis allows researchers to integrate sedimentological, ecological, morphological, taxonomic, and any other data retrieved from geohistorical records. This approach ultimately allows an integrated investigation of the complex interactions between plate tectonics, climate, and the evolution of life.

## 2. The Map Equation framework for network analysis

Network scientists have developed various methods for different purposes and research questions. The unsupervised learning task known as community detection refers to the partition of a network into essential building communities or modules. A particular assignment of the nodes into such modules is called a network partition. Different approaches have emerged from different research goals and motivations, leading to different perspectives on how to formulate the question of community detection on networks (Schaub et al. 2017). The Map Equation framework is an approach for community detection that models random walks on networks and identify the groups of nodes that confine the random-walk flows for a relatively long time (Rosvall and Bergstrom 2008). This dynamic approach provides an intuitive notion of flow-based modules (Smiljanić et al. 2023). In paleobiology research, random-walk flows on networks constructed from geohistorical data capture movement dynamics across nodes beyond the nearest neighbors, such as species moving across their geographic ranges and through geological time. Network-based research in modern paleobiology has largely focused on community detection in networks. Overall, network partitions obtained via community detection with The Map Equation framework have revealed meaningful biostratigraphic (Viglietti et al. 2022), biogeographic (Rojas et al. 2017; Kocsis et al. 2018), and evolutionary patterns (Rojas et al. 2022) (Figure 1A-D).

**Figure 1.**
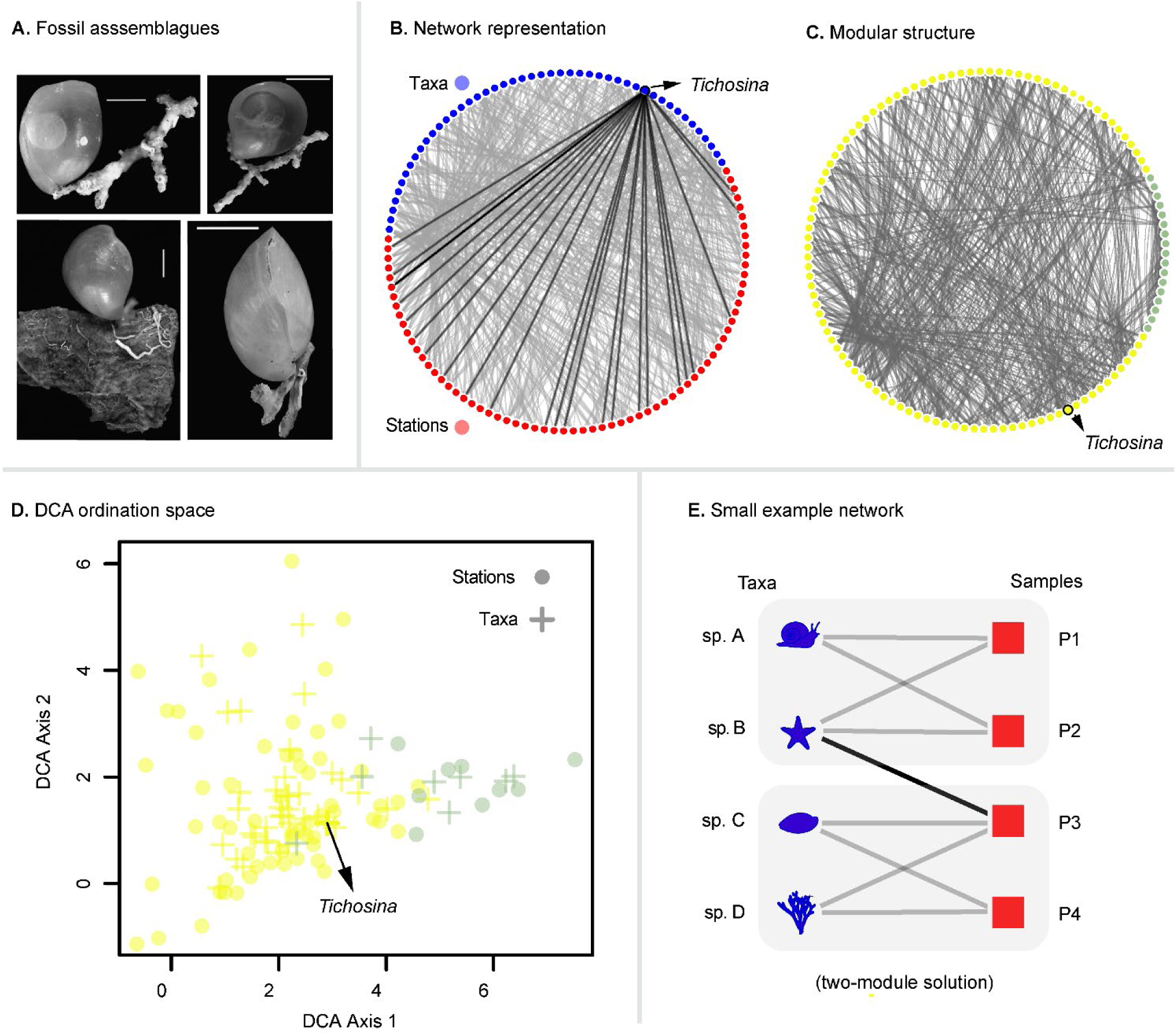
Network representations of geohistorical data. **A.** The physical fossil record. These brachiopod shells aim to represent the benthic macrofauna from the R/V Pillsbury program in the Caribbean. Modified from Rojas et al. (2022, fig. 2). **B**. A network representation of the physical record. This bipartite network is an abstracted representation of the underlying data (sampling stations × taxa matrix) (Rojas et al. 2015)(Supplementary Data 1). The brachiopod *Tichosina*, the larger component of the Cenozoic brachiopod faunas in the Caribbean, is indicated. **C.** Modular structure delineated via community detection with the Map Equation framework and using a Markov Time = 2 (Kheirkhahzadeh et al. 2016) (Supplementary Data 2). Modules include sampling stations and taxa. Nodes are rearranged in the circular layout by their module affiliation. Only the two larger modules, representing 98% of the network flow (see Table 1), are displayed. **D**. Modules mapped on a Detrended Correspondence Analysis (DCA) ordination space. E. Small example showing a bipartite network (1B) partitioned into two modules.

**Table 1.**
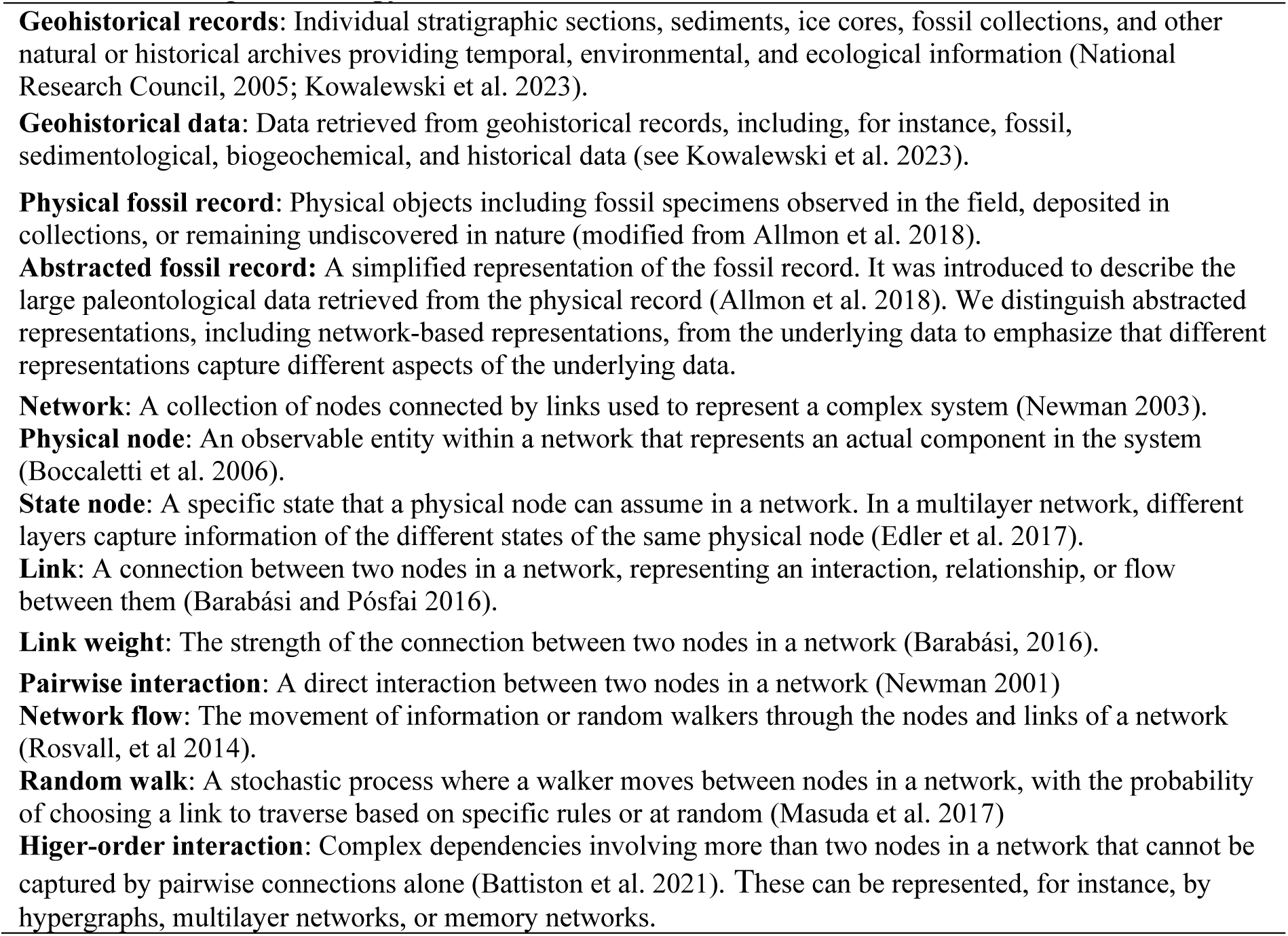
Glossary of terms related to the Map Equation framework in the context of network-based paleobiology research.

The Map Equation framework consists of an objective function – the map equation – and its search algorithm Infomap. The map equation measures the quality of a network partition through the modular description length of the random-walk flows (Rosvall and Bergstrom 2008). Minimizing the map equation over possible solutions with the search algorithm Infomap identifies how network flows organize into modules. Previous research has found that the Map Equation framework performs effectively compared with other approaches on various benchmark networks (Lancichinetti and Fortunato 2009; Aldecoa and Marín 2013; Kheirkhahzadeh et al. 2016; Ghasemian et al. 2019).

### 2.1. The map equation objective function

The map equation is an information-theoretic objective function designed to identify flow-based modules in networks (Rosvall and Bergstrom 2008). It applies the minimum description length principle (Rissanen 1978) to network flows, measuring how well a network partition can compress these flows by capturing their modular patterns. By default, it models network flows as a Markovian diffusion process; a random walker moves between nodes, following links proportional to their weights (Rosvall et al. 2009). For example, in a schematic network representing fossil occurrences across several samples, the random walk begins at a randomly selected sample (Figure 1E), moves to a random taxon present in that sample, and then continues to a random sample where the taxon occurs, and so on repeatedly. When a network contains communities of highly interconnected samples and taxa – representing, for instance, biofacies (Scarponi et al. 2022) or biozones (Viglietti et al. 2022) – the network flows will persist relatively long within those communities.

The map equation captures the network’s modular structure through the stationary node and link visit rates, reflecting the long-term behavior of the random walk (Rosvall and Bergstrom 2008; Rosvall et al. 2009). Given a partition of nodes into modules, it specifies the theoretical lower bound on how concisely we can describe the trajectory of the random walk. An illustration with codewords exemplifies the machinery: A sender wants to convey the random walk’s position after each step to a receiver. Just as Morse code assigns shorter codewords to frequently used letters for efficient communication, the sender uses shorter codewords to frequently visited nodes. For a modular description that capitalizes on the community structure, the sender assigns each node a unique codeword within its module and reuses short codewords across different modules. To ensure the code is uniquely decodable when switching modules, the sender assigns an exit codeword to each module codebook and uses an index codebook with codewords for entering each module.

When the random walker stays within a module, the sender must transmit only a single codeword for the new node. However, if the walker moves between modules, the sender must transmit three codewords: the exit codeword from the current module codebook, the enter codeword for the new module from the index codebook, and the codeword for the new node from the new module codebook. This modular coding scheme becomes efficient when the network partition aligns with the community structure such that the sender can take advantage of shorter codewords within modules and module exits are relatively rare.

The map equation quantifies this tradeoff between efficient descriptions within many small modules at the cost of many transitions between modules and longer descriptions within a few larger modules with rare transitions between them. While we use codewords to illustrate the machinery of the map equation, it exploits Shannon’s source coding theorem (Shannon 1948) and uses Shannon entropy terms to set the lower bounds on the per-step codelengths for the index and module codebooks. For the distribution of visit probabilities *P*, the Shannon entropy *H*(*P*) = −∑*p*_*a*_*log*_2_*p*_*a*_ measures the per-step average codelength in bits. The entropy term for the index codebook is *H*(*Q*) with normalized enter rates *q*_*i*_/*q* into each module *i*, where 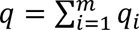 is the total rate at which the index codebook is used. The entropy term for each module *i* is *H*(*P*^*i*^) with normalized visit rates for each node in the module and its exit rate, and *p*_*i*_ is the rate of use for the module codebook. With the entropy terms weighted by their use, the map equation for partition *M* is

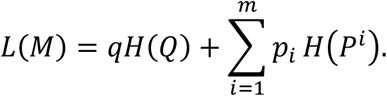

Sample-based geohistorical data typically form undirected networks because the relationship between sampling units and taxa is symmetric: a random walker can move along links between samples and taxa in both directions. In an undirected network, node *a*‘s visit rate is the total weight *w*_*a*_ of its links divided by the total link weight of all *N* nodes in the network, 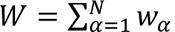. In the small example in Figure 1E, we treat all fossil occurrences as equally important, resulting in an unweighted network where all link weights are equal to 1. Therefore, node *a*’s visit rate is its number of links divided by two times the total number of links in the network, since each undirected link has two link-ends. Similarly, a module’s exit and enter rates are the total number of links that crosses its boundary divided by *W*. In the small example network in Figure 1E, one link with weight 1 crosses each boundary such that all enter and exit rates are 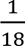. With a compressed notation for the entropy, 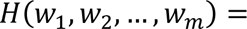 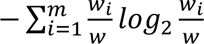, where 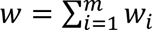, the total codelength for the two-module partition is

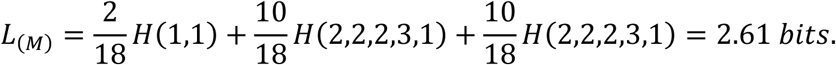

This partition minimizes the map equation (Figure 2). In general, small modules where random walks persist long compress the network flows maximally and reveal the most modular regularities for the modeled network flows (Rosvall and Bergstrom 2008; Rosvall et al. 2009).

**Figure 2.**
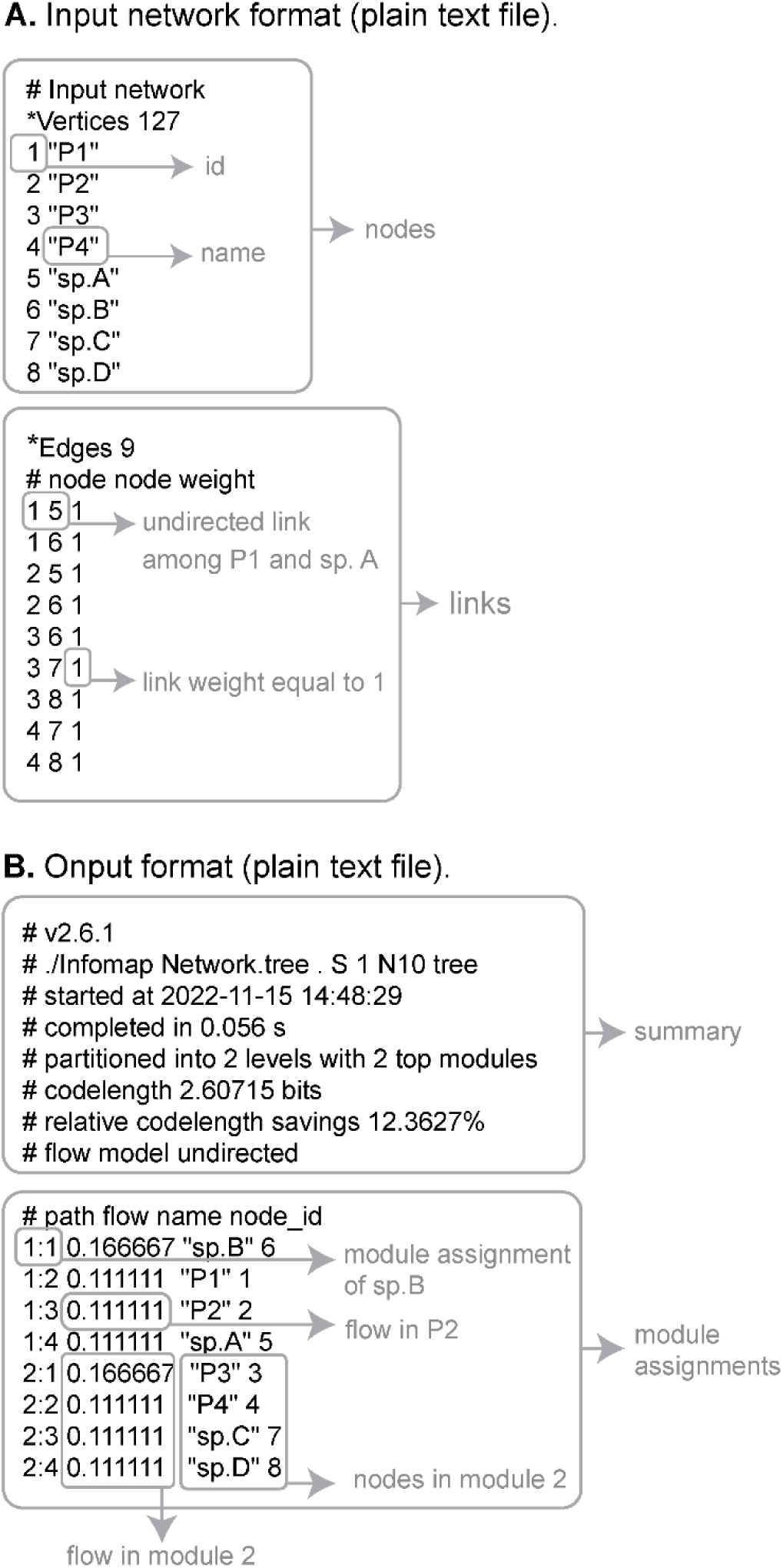
Input and output formats. -The Map Equation framework understands different formats. A. Input file in Pajek format. B. Output file. The resulting partition is written to a file with the extension .tree (plain text file). The formats described here correspond to the small example network in Figure 1E.

The map equation generalizes straightforwardly to multilevel, hierarchically nested modules (Rosvall and Bergstrom 2011), to higher-order network models, including so-called memory networks (Rosvall et al. 2014), multilayer networks (De Domenico et al. 2015), and hypergraphs (Eriksson et al. 2021, 2022), and to varying Markov time models (Kheirkhahzadeh et al. 2016). Recent Bayesian generalizations of the Map Equation deal with incomplete data in sparse networks (Smiljanić et al. 2020, 2021).

### 2.2. The search algorithm Infomap

Infomap is a greedy stochastic search algorithm designed to minimize the map equation over possible node assignments and detect two-level and multilevel flow modules in networks (Rosvall and Bergstrom 2008). Since community detection in networks is a complex optimization problem, no search algorithm can guarantee finding the map equation’s global minimum, and it is often informative to investigate a set of good partitions (Calatayud et al 2019). For a network with *n* nodes assigned to *m* modules, there are on the order of *n^m^* possible partitions. Even for moderately sized networks, testing all possible partitions to guarantee that we have found the best one is impractical. Instead, Infomap uses multiple iterative and recursive optimization heuristics to avoid local minima and provide good network partitions (Edler et al. 2017). The core algorithm starts by assigning each node to its own module. Then it repeatedly loops through each node in random order and moves it to the module that reduces the codelength the most. Infomap repeats this procedure until no move decreases the codelength, rebuilds the network with the modules forming nodes at a coarser level, moves these nodes into even coarser modules, and so on until no move reduces the codelength further.

To improve this two-level solution, Infomap alternates between a fine-tuning and a coarse-tuning procedure by moving individual nodes or sub-modules between modules. To find a hierarchical solution, Infomap starts from the two-level solution and iteratively builds super-module levels that compress the description of movements between modules. Then it clears the structure under each of the coarsest modules and recursively and in parallel builds sub-modules within each module until it cannot find a finer structure that decreases the hierarchical codelength. In this way, the resulting hierarchical structure of the network may have branches of different depths. Tests comparing various community detection methods have consistently demonstrated that Infomap performs exceptionally well across a range of benchmark networks and real-world case studies (Lancichinetti and Fortunato 2009; Aldecoa and Marín 2013; Kheirkhahzadeh et al. 2016; Ghasemian et al. 2019).

### 2.3. The Map Equation for higher-order networks

The Map Equation framework extends to higher-order networks, offering a powerful approach for modeling complex systems with diverse link types and memory effects (Rosvall et al. 2024, Edler et al. 2017). When modeling network dynamics on standard networks through random walks, the nodes simultaneously represent the physical components of the system – observable entities such as individual species, collections, geological outcrops, or stratigraphic units of any resolution, from individual beds to entire formations – and define the flows through pairwise links between them (Figure 3A-B). To overcome the limitations of analyzing network dynamics based solely on pairwise interactions, the map equation for higher-order networks uses so-called state nodes to capture higher-order dependencies and extends Markovian dynamics from first to higher orders (Smiljanić et al. 2023). Network representations with state nodes are referred to as memory networks when these nodes retain information from previously visited physical nodes. State nodes can also reflect various aspects of the system, such as distinct layers in multilayer networks.

**Figure 3.**
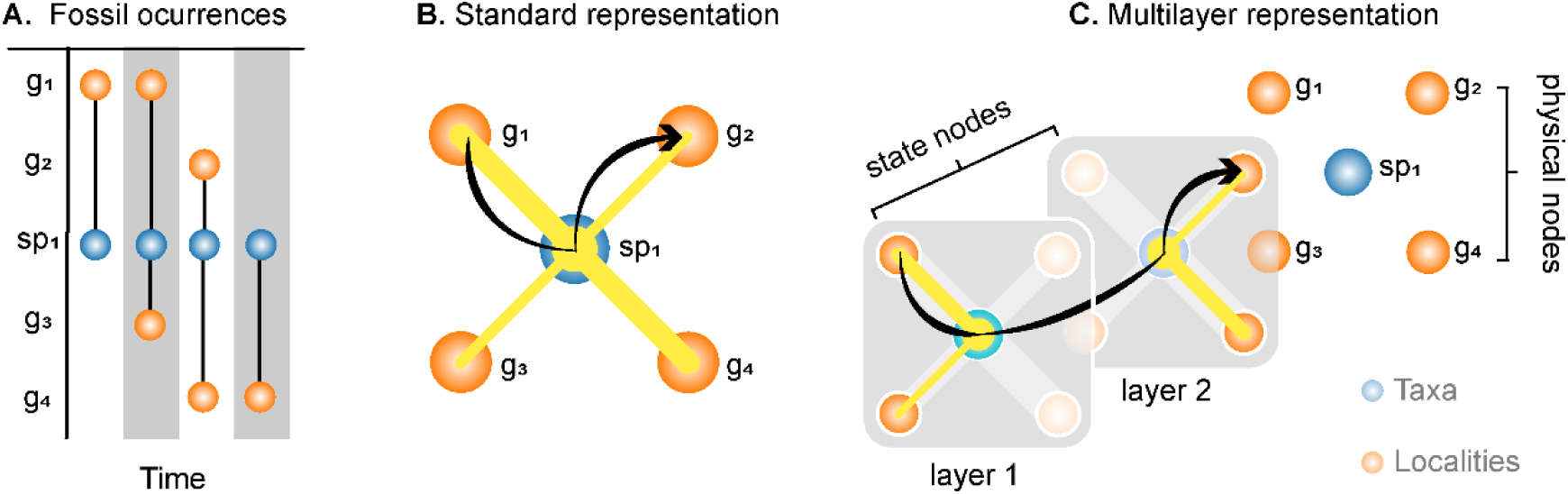
Network representations of geohistorical data. A. Temporal occurrence data. B. Standard bipartite network representation created by aggregating fossil occurrences (sp. 1) into arbitrary spatiotemporally explicit units (g1 to g4). This standard network uses the same nodes to represent the physical components of the system as well as to describe system’s flows (see Table 1). A random walk as the simplest information flow model forms paths across independent pairwise links on this network washing out higher-order node interactions. C. Multilayer network representation. The Map Equation framework for higher-order networks distinguishes physical nodes that represent the system’s components from state nodes describing the system’s internal flows. This higher-order network better captures the constraints on the information paths and thus flows tend to stay among units from each layer.

State nodes not only provide a more accurate representation of real dynamics but also enable overlapping modules, as state nodes of the same physical node can be assigned to different modules. Infomap applies the same iterative and recursive optimization procedures to the state nodes as to standard nodes. However, to generalize the coding machinery to higher-order dynamics and encode only visits to physical nodes, it aggregates the visit rates for all state nodes of the same physical node assigned to the same module, representing them with a single codeword (Edler et al. 2017). This approach has been used to study the Phanerozoic fossil record using a multilayer network representation of global occurrence data, with physical nodes representing interacting taxa and grid cells (aggregated data), and layers representing ordered geological stages. In this case, state nodes denoted the different geological stages where a particular taxon occurs (Rojas et al. 2021). While all intra-layer and inter-layer links can be explicitly provided, when inter-layer links are missing, as in this example, Infomap can generate them through inter-layer relaxation (Figure 3). Since an individual taxon exists in several geological stages, a physical node contains as many state nodes as geological stages where the taxon occurs. As a result, Infomap can provide an optimized solution with taxonomically overlapping communities.

## 3. Higher-order network models in paleobiology

### 3.1. The complex relational structure of the geohistorical data

Geohistorical records, either stratigraphic sections, boreholes, ice cores, or archaeological sites, are inherently complex. Despite their limitations (Kidwell and Holland 2002; Kidwell and Tomasovych 2013), the high-dimensional and spatiotemporally resolved data retrieved from individual geohistorical records allow for evaluation of past biotic responses to natural and human-induced environmental changes at local to regional scales (Council 2005; Scarponi and Kowalewski 2007; Dietl and Flessa 2011; Durham and Dietl 2015; Kowalewski et al. 2023). Although high-precision chronological studies of selected stratigraphic sections have improved our understanding of major biotic crises in Earth’s history (Smith et al. 2018), extensive compilations of data from individual geohistorical records are required for studies at large spatiotemporal scales. Fossil occurrences of the benthic marine invertebrates in the Paleobiology Database (PaleoDB)(Peters and McClennen 2016) have become the benchmark data for network-based research on macroevolution, macroecology, and pleobiogeography (Rojas et al. 2017, 2021; Kocsis et al. 2018; Muscente et al. 2018). In most cases, PaleoDB collections have geographic information and are assigned to a geological stage, enabling the modeling of temporal constraints. PaleoDB collections also have lithostratigraphic and sedimentological information and sometimes include taphonomy and body-size data. Each fossil occurrence in PaleoDB belongs to one collection, has a name with a specific taxonomic resolution, and is linked to an independent taxonomic classification with associated ecological information. These complex relational data describe the structure of the Phanerozoic life at multiple taxonomic levels, stratigraphic resolutions, and spatiotemporal scales.

There are also numerous databases covering specific taxonomic groups, time intervals, or geographic regions (Williams et al. 2018). For instance, The Strategic Environmental Archaeology Database compiles high-resolution archaeological and paleontological data (Buckland and Eriksson 2014). In most cases, the sample’s age is a value taken from the original literature sources and obtained from a range of dating methods varying in precision and accuracy (Buckland 2014). Because samples may have an age range larger than the length of the preferred bin interval, modeling temporal constraints of high-resolution data using multilayer networks is challenging. This question has been approached using the Map Equation framework to investigate the recent fossil record of European beetles (Pilotto et al. 2022). Their multilayer network analysis relaxes the temporal constraints, allowing a random walker to move toward neighboring layers without exceeding the age limits of the samples in the data. In practice, with ordered layers representing 500-year time intervals and an accepted age range of 2000 years in the filtered samples, authors allowed a random walker to relax toward the first two layers in each direction (i.e., *relax limit* = 2) to explore the temporal connections between species and geological sites only within the maximum age uncertainty. This procedure can be modified to allow a variable Relax limit that constrains the random walk movement to the given number of neighboring layers within the age limits of the geological site where it is coming from. Overall, this approach accounts for the age uncertainty inherent to the samples, making it possible to explore high-resolution geohistorical data using multilayer representations.

### 3.2. Higher-order networks capture the complexity of the geohistorical data

One of the major conceptual changes in modern paleobiology research has been the distinction between the physical fossil record, consisting of *in-situ* or *ex-situ* specimens, and abstracted representations based on data retrieved from this physical record (Sepkoski 2013; Allmon et al. 2018). We capitalize on this idea by explicitly considering network representations of geohistorical data as abstracted records that can be improved (Figure 4). Intuitively, a good model of the physical fossil record should be maximally parsimonious yet sufficiently complex to capture the complex interactions of spatiotemporally resolved and high-dimensional geohistorical data. Researchers designing network representations of geohistorical data must recognize that their choices make various assumptions about the spatiotemporal structure and dynamics of the fossil record, such as whether or not to describe temporal constraints. These assumptions impact the outcome, such as whether or not the solutions capture larger-scale patterns. In paleobiology, researchers tacitly assume that working on the same data (e.g., global-scale records of the benthic marine invertebrates from the PaleoDB) guarantees reproducibility, ignoring that different network representations and modeling decisions may impact the observed patterns. Here we argue that setting benchmarks is required to improve reproducibility and communicability in the emergent field of network paleobiology. This challenge is also an opportunity to move beyond standard network representations, toward higher-order models (Battiston et al. 2020) assuming biotic and abiotic components of the environment can interact in higher-order combinations, for instance, the interactions between two species can be affected by other species (Bairey et al. 2016). Overall, these higher-order network models allow to capture the complexity of the biosphere over evolutionary time scales.

**Figure 4.**
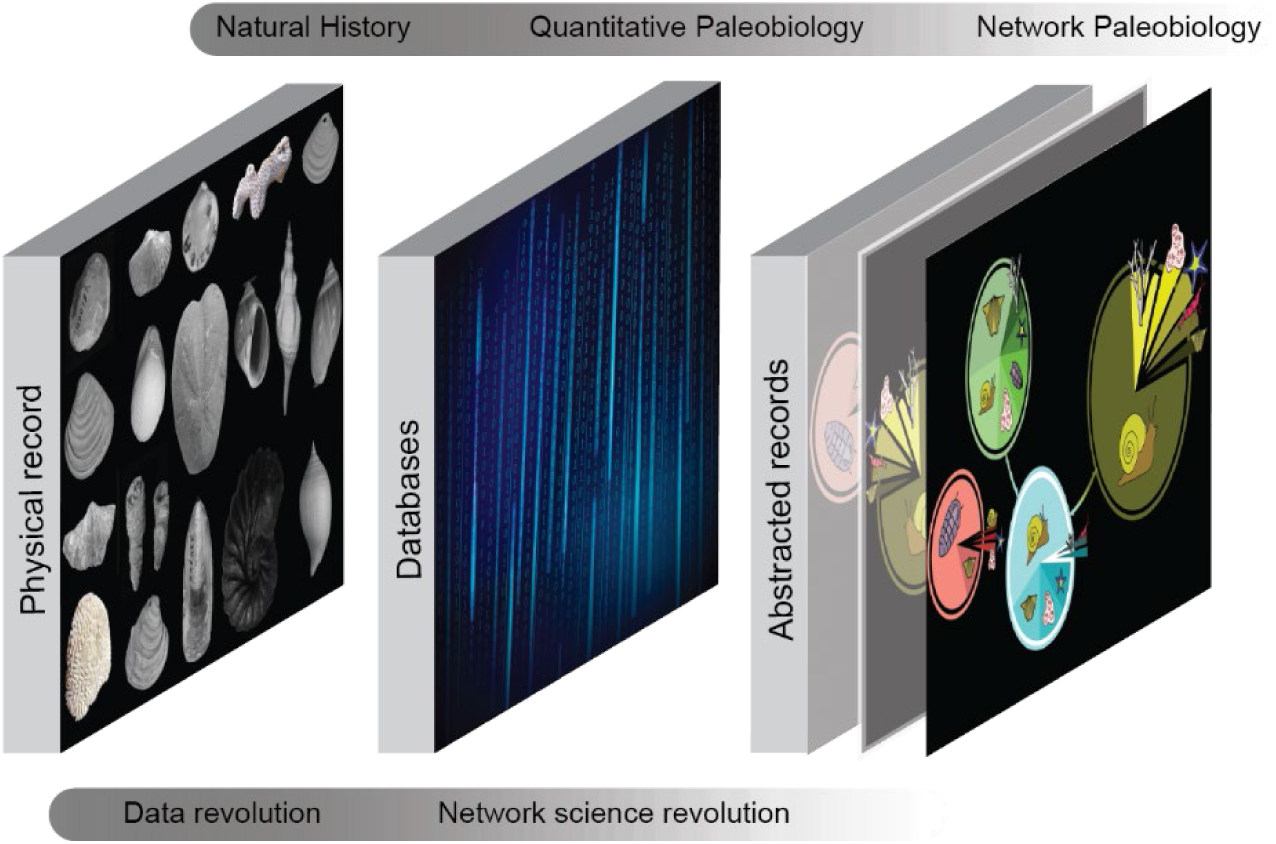
The notion of multiple fossil records in modern paleobiology research. A major conceptual change in modern paleobiology research has been the distinction between the physical fossil record, consisting of specimens, and the abstracted fossil record derived from this physical record (Allmon et al. 2018). The concept of the abstracted fossil record was first introduced to highlight the extensive data already stored in paleontological databases (Sepkoski 2013). This major shift in paleobiology research represents a transition from a specimen-based discipline to its current status as a data-driven field (Dillon et al. 2023). Here we argue that a new major conceptual change in paleobiology has emerged from the growing body of network-based studies that consider network representations of fossil data as abstracted records. From a given physical fossil record and associated data, paleobiologists can build different abstract records that capture different aspects of the underlying system.

Network representations of geohistorical data in modern paleontological literature are typically models accounting for observed interactions among pairs of taxa (e.g., Muscente et al. 2018), sampling units (e.g., Kocsis et al. 2018), or pairs comprising both taxa and sampling units (e.g., Swain et al. 2022) (Figure 3A). This standard approach models dynamical processes on these networks using only pairwise interactions, ignoring a fundamental feature of any complex system, that is the rich pattern of higher-order dependences between components (Battiston et al. 2020) (Figure 3B). In the Map Equation framework, such dependencies are modeled by equipping a random walker with memory, and letting it choose its steps based on its current node but also on the previously visited node or nodes. Whereas standard network models, which actually suffers from memory loss, use a single node type to represent the physical components of the system, including for instance taxa and sampling units, and model flows through one-step dynamics on their links, the Map Equation framework for higher-order networks introduces abstract nodes to describe the different states in which the physical nodes can be in the system, so-called state nodes (De Domenico et al. 2015) (Figure 3C). For instance, the multilayer network representation of the benchmark data on the Phanerozoic fossil record is a higher-order model in which a physical node representing a given taxon contains several state nodes carrying information on the geological stage where this taxon occurs (Rojas et al. 2021). Therefore, this higher-order representation created through the Map Equation framework is a form of sparse memory network (Edler et al. 2017). In addition to the information inherent to the species, states nodes can be used to describe the localities or any other sampling units representing physical nodes in the network, for instance, they can be used to indicate the tectonic context of the grid cells resulting from aggregating global occurrences (i.e., a given grid cell will have as many state nodes as the number of different plates it lies on over time). Overall, first-order models of stage-level occurrence data of the Phanerozoic benthic marine faunas without memory ignore some of the constraints inherent to the underlying system (e.g., the law of faunal succession, geohistorical time arrow, place tectonic context, etc.) and may obscure the macroevolutionary patterns (Figure 5). Furthermore, when comparing optimized partitions of unipartite, bipartite, and multilayer representations of this data, the multilayer network achieves the shortest codelength and the best compression (see below; Table 2).

**Figure 5.**
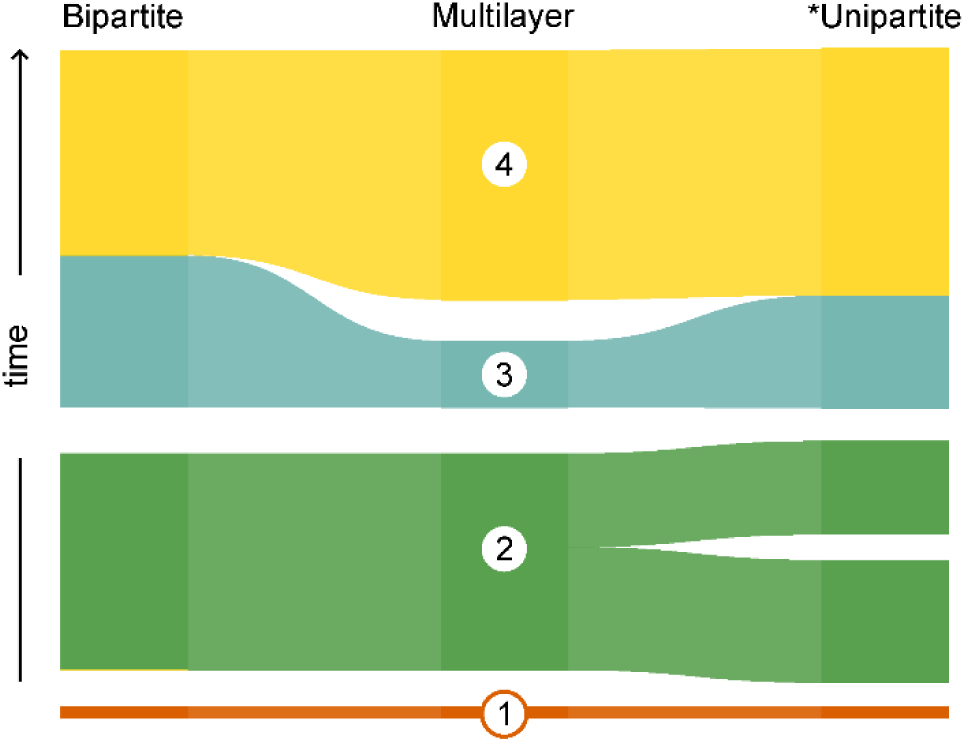
Different network representations capture different aspects of the benchmark data on the Phanerozoic benthic marine faunas (Peters and McClennen 2016). Bipartite and unipartite representations ignore the temporal constraints inherent to the biosedimentary record. *Unipartite projection obtained from the bipartite network by rescaling the Markov time (Kheirkhahzadeh et al. 2016). Alluvial diagram representing 98% of the network flow in each case (Supplementary Data 3-7).

**Table 2.** Comparison of the modular structure of different network representations of the benchmark data on the Phanerozoic marine faunas. The multilayer representation achieves the shortest codelength and the best compression. For each partition, we measure the compression by using the corresponding one-level partition as a baseline. *Unipartite projection obtained from the bipartite network by rescaling the Markov time (Kheirkhahzadeh et al. 2016).

| Model | Physical nodes | State nodes | Levels | Top modules (99%) | Codelength (bits) | Compression (%) |
| --- | --- | --- | --- | --- | --- | --- |
| Unipartite* | 23203 | — | 2 | 4 | 11.0646 | 8.6009 |
| Bipartite | 23203 | — | 3 | 3 | 10.5169 | 13.1253 |
| Multilayer | 23203 | 64508 | 7 | 4 | **6.2826** | **55.2804** |

### 3.3 Higher-order network representations for geohistorical data

#### 3.3.1 Multilayer networks

A multilayer network is a model used to represent a system with multitype interactions between its components. In the Map Equation framework, multilayer networks can be used to model temporal and non-temporal constraints in the data. Similar to the geological principle of superposition used to determine the relative ages of layered strata, layers in multilayers can be used to capture the ordered arrangement of rock (lithostratigraphic) and time-rock (chronostratigraphic) units (Rojas et al. 2021; Viglietti et al. 2022), (Viglietti et al. 2022)as well as equal-interval time bins (Pilotto et al. 2022). Interlayer dynamics in multilayer networks is often modeled based on the intralayer information (Eriksson et al. 2022). For instance, with layers representing geological stages, the intralayer link structure can describe species occurrences at the localities in each stage, whereas interlayer connections are modeled linking the same species across stages. This is achieved through neighborhood flow coupling (Aslak et al., 2018) between state nodes of the same physical node (or observable entity), representing the same species in different layers (see Figure 3). In practice, the Map Equation framework can generate interlayer links using a *relax rate* (*r*), with a random walker moving between nodes within a given layer guided by intralayer links with probability 1−*r* and relaxing to other layers guided by links between state nodes of the same physical node with a probability *r* (Edler et al. 2017). Previous studies show that a relax rate equal to 0.25 is large enough to capture interlayer structures but small enough to preserve intralayer information (Aslak et al. 2018). This feature is useful when measuring interlayer links empirically is challenging. By gradually tuning the *relax rate*, it is possible to explore the relative contribution of intra- and interlayer connections to the overall network structure (Farage et al. 2021).

Multilayer networks can be also used to model multitype interactions from geohistorical data lacking temporal structure o when the research design does not account for changes over time. In general, geohistorical records, including stratigraphic sections, sediments, fossil collections, and ice cores (National Research Council 2005), consist of multiple biotic and abiotic components and can be conceptualized as systems with multilayered structure, where each layer describes a particular interaction between the observable entities. For example, samples from different well sections across a marine basin can be linked together based on their (*i*) fossil content and (*ii*) lithological characteristics, independently. A two-layered network can be used to represent this system, with each layer representing a different type of relationship between the samples. As a consequence, in this particular multilayer network, each observable entity, which is represented by a unique physical node, comprises a number of state nodes equal to the number of interactions in the system, Our first example illustrates multilayer networks through a case study based on the marine invertebrate fossil record (Holland and Patzkowsky 2004).

#### 3.3.2 Varying Markov time models for higher-order networks

The fossil record does not have a unique and optimal level of description but instead multiple levels representing different scales in the organization of life. The Markov time represents the scale of time over which a Markov process evolves on a network. In the context of flow-based community detection such as the Map Equation Framework, the Markov time indicates how information or a random walker propagates through the network over different time scales. Modeling network dynamics at shorter or longer Markov times captures the modular structure at different resolutions (Delvenne et al. 2010). These Markov time models can be used to explore structure and dynamics in both first- and higher-order networks (Kheirkhahzadeh et al. 2016). They are particularly useful when the modular structure of an empirical network lacks a hierarchical organization (e.g., Penn-Clarke and Harper 2020) or when a two-level solution (i.e., non-hierarchical) is used in the Map Equation framework. Exploring various time scales to reveal finer or coarser partitions allows connecting time scales of the dynamics to the structural scales in the network (Lambiotte et al. 2014). Although network analyses in paleobiology typically assume one-step dynamics on the links, corresponding to Markov time 1, a recent study used varying Markov time models to compare the organization of a Holocene succession with present-day nearshore seabed of the Adriatic Sea (Scarponi et al. 2022). However, the specific relationship between Markov time and the multiple time scales of the evolution of the Earth-Life system, recorded in the deep-time fossil record (Rojas et al. 2021), has not been explored.

In the Map Equation framework, the discrete (Kheirkhahzadeh et al. 2016) and continuous (Schaub et al. 2012) time evolution of a Markov process on the network is defined by the parameter Markov time. Intuitively, when using shorter Markov times than 1, the average transition rate of a random walk is lower than the encoding rate (see Map Equation section), the same node is encoded multiple times in its trajectory, and smaller modules are delineated. In contrast, when using longer Markov times than 1, the average transition rate is higher than the encoding rate, only some nodes on its trajectory are encoded, and larger modules are delineated. In practice, by setting *markov time* = 2, we explicitly explore the two-step dynamics on the network. Our second example illustrates varying Markov time models through an empirical study on the mid-Devonian biogeography of the brachiopods where one-step dynamics on the links (*markov time* = 1) in a relatively small bipartite network does not capture a hierarchical organization.

#### 3.3.3 Hypergraphs

Hypergraphs are network models used to study complex systems in which an arbitrary number of its components can interact. These so-called multigroup interactions, represented through hyperedges connecting all the nodes involved in the interaction, differ from binary contacts represented through links in conventional network models (Carletti and Fanelli 2022). A recent study provides the first hypergraph representation of data derived from the fossil record (Eriksson et al. 2021). This network analysis employs the Map Equation framework to model global occurrences of the benthic marine animals from Cambrian (541 MY) to Cretaceous (66 MY), sourced from the PaleoDB, as a hypergraph where physical entities are fossil taxa linked through weighted hyperedges that connect all taxa occurring at each stage. Because this network explicitly models temporal constraints in the underlying paleontological data as hyperedges, it captures large-scale temporal structure and dynamics of system, especially when spatial data are lacking (Figure 6).

**Figure 6.**
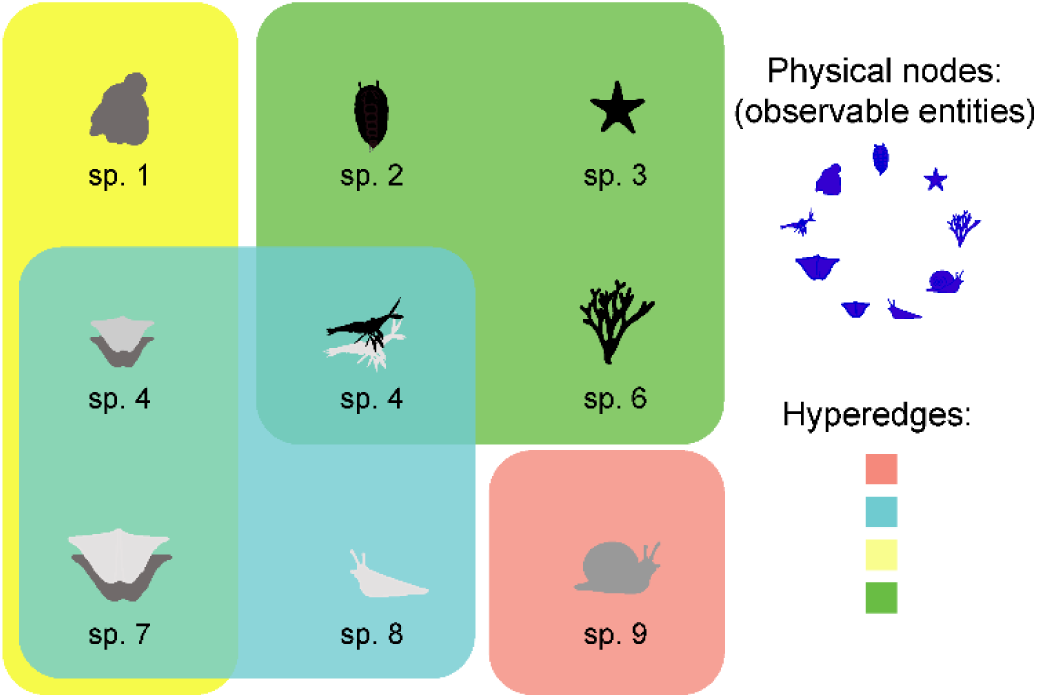
Schematic diagram illustrating a hypergraph representation of fossil occurrence data. In this diagram, observable entities are fossil taxa, represented by shapes of different sizes, connected through hyperlinks. Hyperlinks are depicted with boxes of different colors connecting all taxa (species 1 to 9) occurring at the same time interval. In the Map Equation framework, occurrences of a given taxa at different time intervals are modeled as state nodes, which are depicted here with different shades of grey.

This recent study illustrates how to use the Map Equation framework to model multigroup interactions in fossil occurrence data (Eriksson et al. 2022). It creates unipartite, bipartite, and multilayer network representations of hypergraph flows and evaluated how the choice of both network representation and specific random-walk model, e.g., non-lazy and lazy random walks, with the latter allowing staying at the current node (or self-linking) with a certain probability, impacts the number, size, depth (i.e., number of hierarchical levels), and overlap of multilevel communities in geohistorical data. To create each different network model, this study represents hyperedges as nodes in the bipartite network, projected the bipartite flow to create a unipartite flow representation (i.e., the two-step dynamics on the network obtained with *markov time* = 2) (Kheirkhahzadeh et al. 2016), and created state nodes for each hyperedge to which a node belongs to construct the multilayer network. Overall, the results illustrate the advantages of using multilayer network representations of data derived from the fossil record over bipartite and unipartite representations to quantify macroevolutionary patterns, with different random walk models, including and excluding self-links, providing similar solutions.

## 4 Visualizing higher-order multiscale structure in networks

Network visualizations are essential tools for understanding the modular structure and dynamics of higher-order networks describing geohistorical data. However, standard graphic tools developed to represent first-order interactions among network components fail to capture higher-order and multiscale community structure and depend on the arbitrary scale of analysis (Peixoto and Rosvall 2017; Perri and Scholtes 2020). To overcome these limitations, the Map Equation framework provides graphic tools for mapping higher-order and multiscale community structure in network partitions. They are freely available as a client-side web application at https://www.mapequation.org.

### 4.1 Infomap Network Navigator (INN)

Higher-order networks representing geohistorical systems are usually complex with hierarchical modular structures. The Network Navigator tool was developed to explore such complex structures in real networks. The tool creates interactive maps of hierarchical network partitions with aggregated inter-module links. These, possibly directed, links are drawn with lengths inversely proportional -- and width and color saturation proportional to the flow volume between modules. Weakly connected modules are placed further apart with narrower links between them than strongly connected modules. Circles represent the modules, with areas proportional to the contained flow volume and border thicknesses proportional to the exiting flow (Figure 7). Like Google Maps, the modules can be explored by zooming in and highlighting more detail until the lowest-level leaf nodes are shown. The Network Navigator was recently used to visualize the multiscale organization of the Phanerozoic benthic marine faunas, highlighting how the large-scale evolutionary faunas are built up from lower-scale biogeographic entities (Rojas et al. 2021). Although the INN uses a force-directed drawing algorithm, similar to those used in the traditional visual approach to draw observed entities and their connections and to gain insights into the network structure (e.g., Huang et al. 2016; Penn-Clarke and Harper 2020; Ye et al. 2021), the INN actually represent the network structure previously delineated via community detection. Nevertheless, given the challenge of visualizing substantially different network partitions using the same approach, the INN incorporates graphic parameters that allow users to scale the module size and link width, and to select the level of detail displayed in the plot, making the visual output easier to interpret in each case.

**Figure 7.**
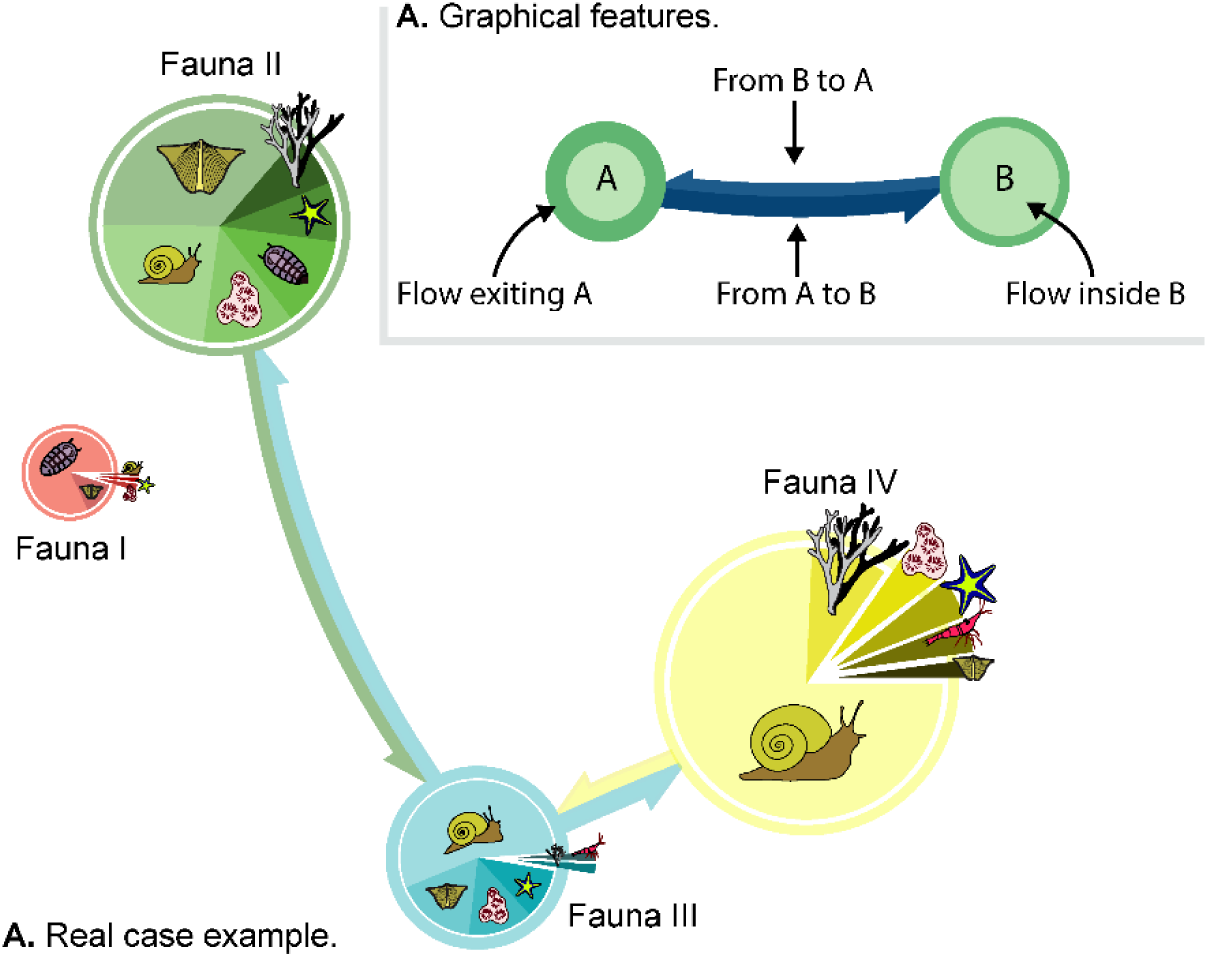
Infomap Network Navigator. **A.** Graphical Features. Modules are represented as circles, with areas proportional to the internal flow and border widths proportional to the exiting flow; aggregated bidirectional inter-module links are shown with widths proportional to inter-module flow and lengths inversely proportional to it. **B**. Real Case Example. Stylized network navigator output illustrating the large-scale modular structure of Phanerozoic benthic marine faunas, based on data from Rojas et al. (2021).

### 4.2 Alluvial diagrams

Researchers compare network partitions typically to identify major changes in the overall modular structure. However, common visualization tools employed in network-based paleobiology research, which focus on the link structure among the observable entities, aim to represent all individual nodes and their connections (Muscente et al. 2019; Penn-Clarke and Harper 2020). This approach is challenging given the high-dimensional data but also typically unnecessary in order to answer common research questions in paleobiology (e.g., to delineate bioregions, biozones, mega-assemblages, biotic transitions, etc.). Alluvial diagrams are visualization tools that highlight changes in modular structure across different network partitions, including, for instance, optimized, bootstrapped, suboptimal, and planted solutions. In the recent paleobiology literature, alluvial diagrams have been used, for instance, to compare a Holocene succession with present-day nearshore seabeds of the Po Adriatic system (Scarponi et al. 2022), and to highlight major differences between alternative biostratigraphy models for the late Permian-mid Triassic Beaufort Group in South Africa (Viglietti et al. 2022). Overall, an alluvial diagram does not represent all individual nodes in the network but highlights changes in the module assignment of the most important nodes across partitions. In the Map Equation framework, the flow volume determines node importance and is calculated from the visitation rates when the Infomap algorithm searches for the optimal partition (Rosvall and Bergstrom 2008).

To visually simplify patterns while highlighting changes between network partitions, the alluvial diagram is constructed by grouping nodes with the same module assignment as follows (Figure 8): (*i*) Shown side-by-side, each partition is represented by a vertical stack of modules in the order we choose. (*ii*) To highlight the nodes that change module assignment between partitions, we draw streamlines between all modules in adjacent partitions that contain the same node. (*iii*) The flow volume of the modules determine their heights, whereas the flow of the nodes that connected modules have in common determine the height of the streamlines. The alluvial diagram comparing network partitions obtained from different network representations of the same underlying data (Figure 5) illustrates how this graphic tool helps to better understand changes in the modular structure between partitions.

**Figure 8.**
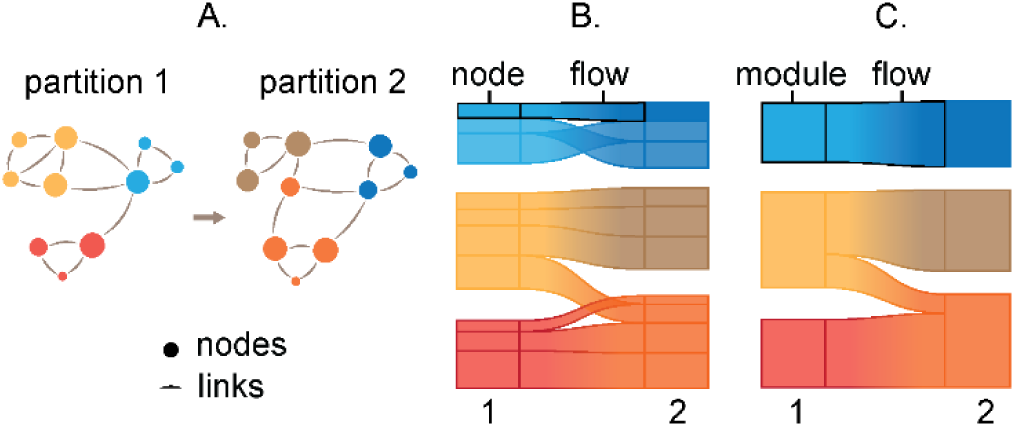
Visualizing change in network partitions using alluvial diagrams. To compare networks with the same sets of nodes, we assemble at least two network partitions. **A**. Then, we group nodes in the same module in stacked bars with height proportional to the flow volume of each node and connect corresponding nodes with streamlines. **B**. Finally, to highlight how the partitions change, we aggregate the nodes into modules. **C**. To compare additional network partitions, we add more stacks of bars to the right and repeat the procedure b-c.

## 4. Robustness evaluation

### 4.1. Identifying significant assignments in network partitions

To assess the support of network modules in the underlying data, it is convenient to construct a set of bootstrapped networks through resampling of the data in a standard manner (Efron and Tibshirani 1993). This approach enables calculating summary statistics such as mean values and standard deviations and identifying features that occur in a large enough proportion of bootstrapped networks to be considered robust. The Map Equation framework includes a significance clustering method (Rosvall and Bergstrom 2010), that identifies sets of nodes that are significantly assigned to a module in a reference partition. For a given module in the reference partition, the largest subset of nodes clustered together in at least a user-defined proportion of all bootstrap partitions represents its significant core. This confidence level is provided as a fraction with the default value being 0.95. This approach has been used to distinguish gradual from abrupt events in the deep-time (Rojas et al. 2021) and near-time fossil record (Pilotto et al. 2022), with gradual events assumed when the significant core of two adjacent modules co-occurs more than a fraction [1 - confidence level] of the bootstrap partitions. In addition, different aspects of a network partition can be evaluated across a set of bootstrap networks using set-theoretic measures such as the Jaccard index and measures built upon concepts from information theory (Vinh et al. 2009). The resampling procedure can also consider the underlying data by, for example, considering a discrete distribution if the data represent counts, and also a truncated distribution to avoid false negatives.

### 4.2. Exploring alternative solutions

Finding the best network partition is generally a non-convex optimization problem and the practitioner therefore needs to consider the possibility of multiple solutions. To this end, methods exploring solutions and their quality have been developed. One such method estimates the minimum number of searches required by the search algorithm Infomap to map the complete solution landscape, ensuring that the best solution is obtained and that alternative solutions of lesser quality can be explored (Calatayud et al. 2019). We illustrate this approach with the multilayer network representing the Phanerozoic fossil record of the benthic marine faunas, where alternative partitions are embedded using a dimension reduction technique (McInnes et al. 2018)(Figure 8), and the distance between partitions is calculated using a weighted version of the Jaccard distance. In this example, the codelength (see Table 1) varies between the partition clusters, with the best partition in the middle cluster. In practice, these variations have minimal impact on the large-scale patterns, with only a few nodes alternating between modules.

**Figure 9.**
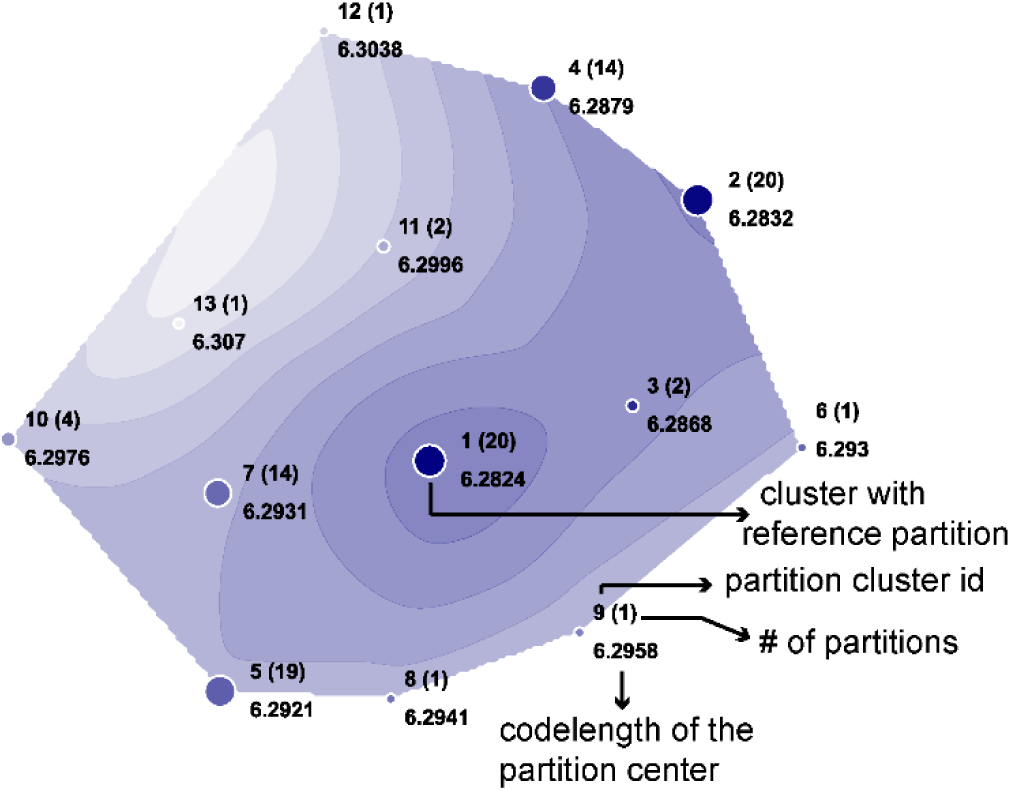
**A**. Quality of alternative partitions of the multilayer network representing the Phanerozoic fossil record of the benthic marine faunas mapped in a two-dimensional space. Circles represent clusters of network partitions, located at the cluster center, with size proportional to the number of partitions grouped into the cluster. Map isolines are constructed using the Jaccard distance between partitions. Despite differences in their quality (codelength), all partition clusters identified at the selected scale show a similar modular structure

Although the approach to explore the solution landscape was developed to identify the minimum number of searches required by the search algorithm Infomap to map the complete solution landscape, it also highlights the importance of exploring sub-optimal partitions when dealing with real systems, which can provide a better understanding of complex patterns. For instance, the end-Cretaceous extinction, despite being a widely known event (Alegret et al. 2022), is not captured at the resolution of the landscape illustrated in Figure 8. To find a solution showing the K-T event at the highest hierarchical level, we must consider less optimal solutions, suggesting that other global events have been more important in shaping the Phanerozoic history of the marine life.

## 5. Case studies on the fossil record

### 5.1. Delineating litho-biofacies through multilayer networks

Understanding how the distribution of organisms along environmental gradients changes through time is a primary research area in paleobiology (Patzkowsky and Holland 2012). Indirect ordination techniques applied to species occurrence data have been successfully employed to recover environmental gradients in the sedimentary record (Holland and Patzkowsky 2007; Dale et al. 2007). In general, environmental gradients are interpreted from mapping taxa or samples, coded by external factors (e.g., life habit for taxa and depositional environment for samples), into the reduced ordination space. Here we provide a multilayer network analysis that combines taxon abundance and sample attributes into the modeling. Specifically, we conceptualize the biosedimentary record as a complex system with a multilayered structure by creating a network representation with two layers, one describing taxonomic composition and the other sedimentologically-defined relationships between sampling units (Figure 10A). The underlying data were obtained from a basin-scale study carried out on Late Ordovician outcrops in central Kentucky (Holland and Patzkowsky 2004)..

**Figure 10.**
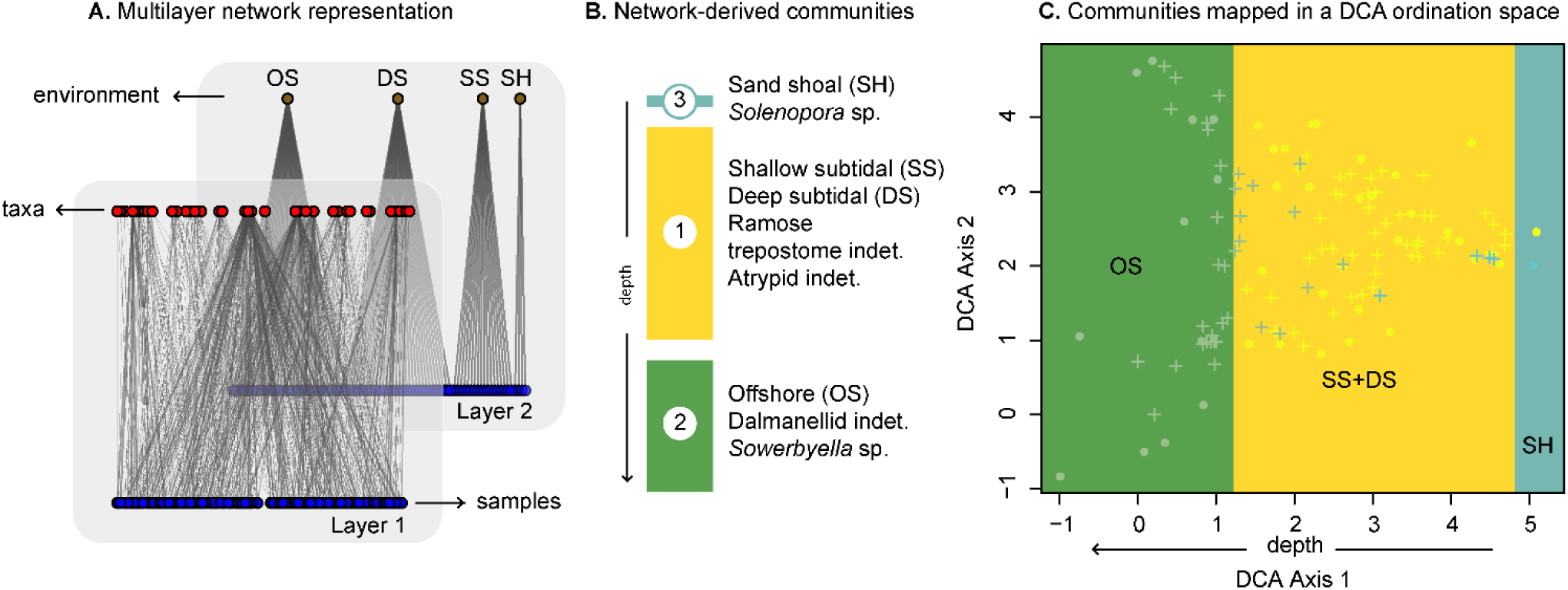
**A**. A multilayer network representing litho-biofacies from Late Ordovician outcrops in central Kentucky. In this higher order network, one layer describes the biotic component (samples × taxa matrix) and the other describes the abiotic component (samples × environments matrix) of the geohistorical record (Supplementary Data 8). **B**. Litho-biofacies delineated via community detection using the Map Equation framework. Modules in the multilayer solution include taxa, samples, and environments, and can be directly interpreted as litho-biofacies (Supplementary Data 9). **C**. Detrended Component Analysis (DCA) on the samples × taxa matrix. Network modules mapped in a DCA ordination space indicating their distribution along the depth water gradient. Background colored based on the module affiliation. Data from Holland and Patzkowsky (2004).

Observable entities in this two-layered network represent sampling units, taxa, and sedimentological groups. Layers in this representation independently capture fossil assemblages (biofacies) and sedimentological groups (lithofacies) in the geohistorical record, independently. However, the multilayer network reveals litho-bio facies (Figure 10B). This higher-order approach directly interprets the gradients obtained via ordination analysis in two ways, delineating modules that comprise strongly connected taxa, samples, and sedimentological groups, and partitioning the gradient into discrete regions (Figure 10C). Although the sedimentological information underlying this case study is relatively simple, our network model can be easily extended to represent geochemical, taphonomic, and any other complex relationship between sampling units, as well as relationships between taxa (i.e., ecology, body size). Even though beds are the fundamental units of both stratigraphy and paleontology (Patzkowsky and Holland 2012), sampling units in this multilayer framework can represent stratigraphic units of any scale. In practice, depending on the stratigraphic resolution of the underlying data, this multilayer network model can be used to capture multitype relationships between samples, beds, members, or broader units

### 5.2. Validating marine bioregionalization through Markov time models

Research employing network-based approaches to describe biogeography in the fossil record is overwhelmingly focused on describing one-step dynamics (Markov time 1) in relatively small bipartite networks derived from the relatively limited fossil data (Penn-Clarke and Harper 2020; Ye et al. 2021), to reveal continental to global scale marine bioregions. Although first-order network representations of fossil occurrences have been shown to capture a biogeographic signal at some geological stages in the Phanerozoic (Rojas et al. 2017; Kocsis et al. 2018), these studies are unable to identify transition zones, provide a single-scale description of the bioregions and obscure larger-scale patterns. Overall, conventional approaches ignore that bioregions do not have a unique level of description but multiple levels reflecting the complex spatiotemporal structuring of the marine biodiversity. Here we describe how to use varying Markov time models through the Map Equation framework to overcome some of the limitations of the standard models currently used in paleobiology research. This case study is based on a standard bipartite network relevant for Devonian biogeography, but the approach can be applied directly to higher-order networks (Calatayud et al. 2021). We use varying Markov time models to re-examine the bioregionalization of the Middle Devonian Brachiopods from the Old-World Realm (Penn-Clarke and Harper 2020). This approach reveals the larger-scale biogeographic structure at different resolutions. Results provide new insights into the biogeographic affinities of the brachiopod faunas from Southern Peru, which remains an open question (Figure 11). However, this case study shows that we can reveal the spatially nested hierarchical organization of marine biodiversity through varying Markov time models.

**Figure 11.**
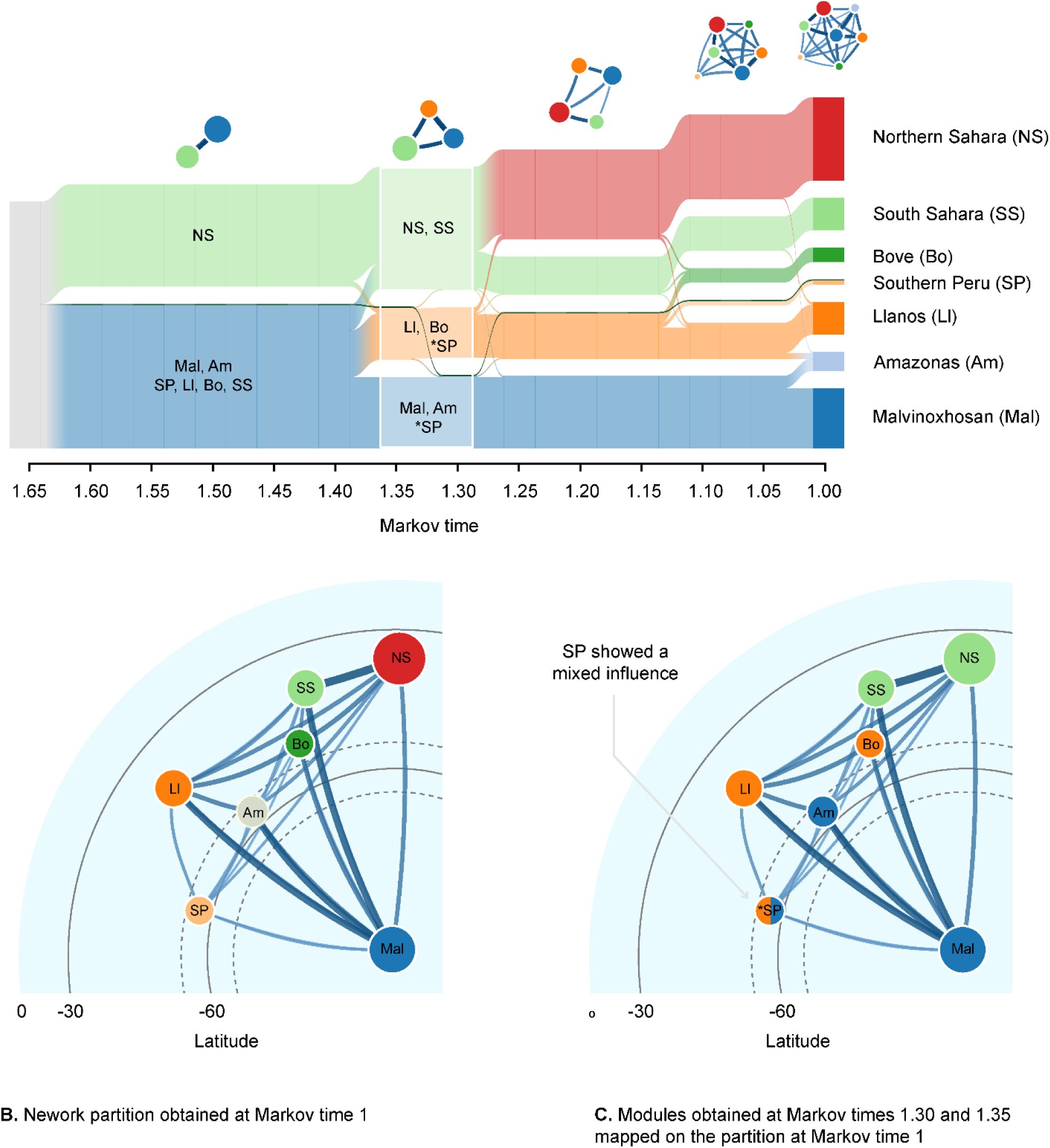
Bioregionalization of the Middle Devonian Old World Realm. **A**. Varying Markov time models on the biogeographic network constructed from brachiopod occurrence data (Penn-Clarke and Harper 2020). Network partitions at different Markov times reveal the larger-scale biogeographic structures obtained at different resolutions. **B.** Network partition obtained at the Markov time 1 (7 modules). Circles represent the seven modules delineated when exploring the one-step dynamics on the assembled network. **C.** Modules obtained from partitions at Markov times 1.30 and 1.35 (3 modules) mapped on those obtained at the Markov time 1. These two partitions differ in the affiliation of the Southern Peru locality, placed alternatively into the modules representing higher (white) and lower (orange) latitudes. Overall, coarser partitions contain fewer clusters as the Markov time increases.

## 6. Recommendations for future research directions

Higher-order network modeling of geohistorical data holds considerable promise for paleobiology research because it provides a framework for revealing the complex interactions between biotic and abiotic components of the geohistorical records at multiple scales. Higher-order networks better capture the spatiotemporal constraints inherent to the geohistorical data, providing more accurate descriptions of patterns. We showed that different network representations capture different aspects of the underlying data and studied systems. This can lead to potential reproducibility issues when researchers overlook how their choice of a network model impacts the results and provide inadequate or incomplete explanations of the analyses (e.g., inadequate description of the input network and incomplete explanations of the clustering approach). We believe that establishing benchmark networks from widely used compendia of paleontological data (e.g., The Paleobiology Database) and using the Map Equation framework can enhance both reproducibility and communicability in network-based paleobiology. Our analyses have focused on regional and global-scale examples, but this framework can be applied to systems at any scale, ranging from individual beds up to the global biosedimentary record.

The standard formulation of the map equation implicitly assumes complete data. It reveals the large-scale structure of a system given the observed data (Ghasemian et al. 2019). This general approach dominates network-based research in paleobiology and has proven useful for describing biogeography, biostratigraphy, and macroevolution. However, geohistorical records are incomplete – not every species that ever lived is preserved, and not all environmental conditions of every region in the globe are recorded in the sediments (Kidwell and Flessa 1996; Council 2005). Sampling also varies due to variations in exposure and collection effort (Holland 2016). The overall effect is an abstracted fossil record partially observed, a network involving links that exist but are not represented in the model because they have not been observed. Distinguishing missing links (false negatives) from non-edges (true negatives) within the unobserved connections, a task known as link prediction, could allow to improve abstracted fossil records where the underlying geohistorical data is highly incomplete. Such missing links could alter conclusions when delineating network community structure and modeling network dynamics (Ghasemian et al. 2019; Blöcker et al. 2022). In cases where we expect data to be highly incomplete, we should use a community-detection approach that can handle this: the Bayesian map equation, also part of the Map Equation framework, models missing data with an empirical Bayes approach (Smiljanić et al. 2021). In addition, using the minimum description length principle (Rissanen 1978) underlying the Map Equation Framework, it is possible to use a modular partition to measure the description length of links (Blöcker et al. 2022). Assuming that links with a shorter description are more likely, we can rank the non-existing links in a network and select the best candidates for false negatives, those links that are most consistent with the network dynamics and thus most likely to exist. In practice, it can be used to improve abstracted fossil records, but it is up to the researchers to examine and interpret those predicted links in the face of the specific question at hand. This approach can be used for stratigraphic placement of isolated samples, refining biostratigraphic models, and improving the overall description of bioregions. We believe these methodological efforts lay the ground for a fertile research direction in the emergent field of network paleobiology.

## 8. Author Contributions

A.B.: conceptualization, data curation, formal analysis, methodology, visualization, writing—original draft, writing, editing; A.E.: visualization, writing; M.N.: visualization, writing, editing; D.E.: writing; C.B.: writing, editing; M.R.: writing, editing, funding.

## Acknowledgments

We thank the contributors to the Paleobiology Database who collected data. We thank Alex Dunhill and Andrej Spiridonov for helpful comments on an early version of the manuscript. AR thanks Steve Holland for providing data to delineate Middle Upper Ordovician biofacies.

## Data Availability Statement

Data for this study are available in the Dryad Digital Repository: https://doi.org/10.5061/dryad.sj3tx967z.

